# Speed-specific optimal contractile conditions of the human soleus muscle from slow to maximum running speed

**DOI:** 10.1101/2023.03.19.533354

**Authors:** Sebastian Bohm, Falk Mersmann, Arno Schroll, Adamantios Arampatzis

## Abstract

The soleus is the main muscle for propulsion during human running but its operating behavior across the spectrum of physiological running speed is currently unknown. This study investigated experimentally the soleus muscle activation patterns and contractile conditions for force generation, power production and efficient work production (i.e. force-length potential, force-velocity potential, power-velocity potential and enthalpy efficiency) at seven running speeds (3.0 m/s to individual maximum). During submaximal running (3.0 to 6.0 m/s), the soleus fascicles shortened close to optimal length and at a velocity close to the efficiency-maximum, two contractile conditions for economical work production. At higher running speeds (7.0 m/s to maximum), the soleus muscle fascicles still operated near optimum length, yet the fascicle shortening velocity increased and shifted towards the optimum for mechanical power production with a simultaneous increase in muscle activation, providing evidence for three cumulative mechanisms to enhance mechanical power production. Using the experimentally-determined force-length-velocity potentials and muscle activation as inputs in a Hill-type muscle model, a reduction in maximum soleus muscle force at speeds ≥7.0 m/s and a continuous increase in maximum mechanical power with speed was predicted. The reduction in soleus maximum force was associated with a reduced force-velocity potential. The increase in maximum power was explained by an enhancement of muscle activation and contractile conditions until 7.0 m/s, yet at the maximal running speed mainly by increased muscle activation.

**Summary statement:** The study provides experimental evidence that the human soleus muscle favors contractile conditions for economical work production during submaximal running and for enhancing mechanical power production during maximal running speed.

## Introduction

During steady-state running, the external mechanical power and work that is needed to move the center of mass of the human body increases with increasing speed (Arampatzis et al., 2000; Cavagna et al., 1976; Kaneko, 1990). Across running speeds, the ankle joint remains the key source of mechanical power and work (Arampatzis et al., 1999; Schache et al., 2011; Winter, 1983). The triceps surae muscles are the main plantar flexors and the monoarticular soleus is the largest muscle in this muscle group (Albracht et al., 2008; Crouzier et al., 2018). Accordingly, previous musculoskeletal modeling studies predicted that the soleus produces the major part of mechanical power and work required for running (Lai et al., 2014; Sasaki and Neptune, 2006). Recent studies reported an active shortening of the soleus muscle fascicles throughout the entire stance phase for running speeds between 2.0 and 5.0 m/s (Bohm et al., 2019; Lai et al., 2015; Rubenson et al., 2012; Swinnen et al., 2022), indicating a continuous contractile work production during stance despite a stretch-shortening cycle of the soleus muscle-tendon unit (MTU). The length and velocity decoupling of the muscle belly from the MTU due to tendon compliance and the decoupling of the muscle fascicles from the muscle belly due to fascicle rotation (Alexander, 1991; Azizi et al., 2008) are the two main mechanisms responsible for the different behavior of the fascicles compared to the MTU during running (Bohm et al., 2021a). Active fascicle shortening during MTU lengthening has also been observed for the human gastrocnemius medialis and lateralis during running (Ishikawa et al., 2007; Lai et al., 2018), for the lateral gastrocnemius in guinea fowls during running (Daley and Biewener, 2003) and for medial and lateral gastrocnemius in goats during trotting (McGuigan et al., 2009).

Recently we provided experimental evidence that during slow running (2.5 m/s) the soleus fascicles shorten under conditions that promote economical work production, i.e. at a length close to the optimum for force generation and at a shortening velocity close to the optimum for enthalpy efficiency (Bohm et al., 2021a; Bohm et al., 2021b). Lai et al. reported an increase in the soleus fascicle shortening velocity and a decrease in fascicle operating length from 2.0 to 5.0 m/s running speed (Lai et al., 2015; Lai et al., 2018), indicating a modulation of the soleus contractile conditions to generate force, power and work with increasing speeds. The increased shortening velocity of the soleus fascicles decrease the muscle force potential according to the force-velocity relationship and consequently increase the active muscle volume per unit force and, thus, the metabolic cost of muscle contraction (Roberts, 2002; Swinnen et al., 2023). During submaximal running the soleus fascicles operate on the ascending part of the force-length curve close to the optimal length (Bohm et al., 2019; Rubenson et al., 2012). A decrease of the fascicle operating length with increasing running speed (Lai et al., 2015; Lai et al., 2018) would result in a decrease of the muscle force potential, thus increasing the active muscle volume for a given muscle force as well (Beck et al., 2022). Furthermore, increases in the fascicle shortening velocity with increasing speed (Lai et al., 2015; Lai et al., 2018) may decrease the enthalpy efficiency of muscle contraction (i.e. the fraction of chemical energy that is converted into mechanical work), which is highest at around 20% of the maximum shortening velocity (Barclay, 2015; Hill, 1939). On the other hand, the increase of fascicle shortening velocity will increase the power-velocity potential, which is greatest at ~30% of the maximum shortening velocity. An increasing muscle power-velocity potential enables the production of higher muscle mechanical power (Lutz and Rome, 1994), which is required for fast running. There are currently no experimental studies on humans investigating the soleus operating fascicle length and velocity during the stance phase at running speeds above 5.0 m/s and therefore the soleus contractile conditions for high and maximum running speeds are unknown. Existing musculoskeletal modeling studies predicted a stretch-shortening cycle of the soleus fascicles during the stance phase of high running speeds (Dorn et al., 2012; Lai et al., 2014), indicating a contractile energy absorption during the MTU lengthening phase and thus a different behavior of the soleus contractile element to produce power and work compared to submaximal running speeds.

The main objective of the current study was to gain a better understanding of how muscle contractile conditions (i.e. force-length, force-velocity and power-velocity potentials and enthalpy efficiency-velocity relationship) and activation may influence the soleus muscle force generation, power production and efficiency from slow to maximum running speeds. By mapping the measured fascicle lengths and velocities during running onto the soleus force-length, force-velocity, power-velocity and efficiency-velocity relationships, we assessed its respective potentials and efficiency from 3.0 m/s to the individual maximum running speed. We adopted a Hill-type muscle model and, by using the experimentally determined operating force-length-velocity potential and muscle activation during running as input variables, investigated the contribution of contractile conditions and activation to the predicted force generation, power and work production of the soleus muscle. We hypothesized a gradual shift from advantageous contractile conditions to produce muscle work economically to conditions favorable to produce maximum power with increasing running speed.

## Material and methods

### Participants and experimental protocol

Fourteen young male adults were included in the present study (body height 179 ± 6 cm, body mass 74 ± 8 kg, age 23 ± 4 years, mean ± standard deviation). The participants had experience in sprinting-related training for several years (e.g. track and field, soccer, basketball) and treadmill running and did not report any history of neuromuscular or skeletal impairments in the six months prior to the recordings. The ethics committee of the university approved the study (HU-KSBF-EK_2022_0003) and the participants gave written informed consent in accordance with the Declaration of Helsinki.

A familiarization session including submaximal and maximal running trials on a treadmill preceded the measurements. During the target measurements, the participants ran on the treadmill at seven different speeds: 3.0 m/s, 4.0 m/s, 5.0 m/s, 6.0 m/s, 7.0 m/s, 8.0 m/s and the individual maximal running speed. In each trial, the treadmill speed was increased and then at least five valid steps were recorded at the target speed (i.e. at steady state running). For the determination of the individual maximum, the treadmill speed was increased from either 8.0 m/s or 8.5 m/s in 0.25 m/s intervals based on the expected maximum speed observed during the familiarization session. Two to three speed increment attempts were then conducted until the participants were not able to run faster (termination or personal feedback). During all trials, the participants were secured by the treadmill’s safety harness. A rest of five minutes was given between the running trials. In a separate part of the measurement, we experimentally determined the soleus force-fascicle length relationship using a dynamometer and an ultrasound device. The order of the two measurement parts was randomized.

### Joint kinematics, fascicle behavior and electromyographic activity

During running on the treadmill (hp/cosmos, 190/65 pulsar^®^ 3p), the ankle and knee joint kinematics of the right leg were captured by a motion capture system (Vicon Motion Systems, Oxford, UK, 250 Hz) using an anatomically-referenced reflective marker setup (greater trochanter, lateral femoral epicondyle and malleolus, fifth metatarsal and tuber calcanei). The consecutive knee joint extension maxima over time were used to determine the gait events of foot touchdown and toe-off (Fellin et al., 2010). Cadence (number of steps per second), stance and swing times, step length and duty factor (fraction of gait cycle on the ground) were determined.

The soleus fascicle behavior was recorded at 146 Hz synchronously to the kinematic data by means of a 6-cm linear array ultrasound probe (Aloka Prosound Alpha 7, UST-5713T, 13.3 MHz, Tokyo, Japan) that was placed on the medial aspect of the soleus muscle belly using a custom neoprene-plastic cast. The fascicle length was post-processed from the ultrasound images using a semi-automatic tracking algorithm (Marzilger et al., 2018) that calculated a representative reference fascicle from multiple muscle fascicle portions automatically identified over the whole field of between the manually tracked aponeuroses. A visual inspection of each image was conducted and manual corrections were made if necessary. The fascicle length from five steps, processed using a second-order low-pass Butterworth filter with 6 Hz cut-off frequency, were analyzed and averaged for each speed and participant. For the maximum running speed of four participants only three steps were available for analysis. The pennation angle was calculated as the angle between the deeper aponeurosis and the reference fascicle. The soleus MTU length changes during running were calculated as the product of ankle joint angle changes and the individual Achilles tendon lever arm (see below) (Lutz and Rome, 1996), while the initial MTU length at neutral ankle joint angle (shank segment perpendicular to foot segment) was based on a regression equation (Hawkins and Hull, 1990). The soleus muscle belly length changes were calculated as the differences of consecutive products of fascicle length and cosine of the pennation angle (Fukunaga et al., 2001). Note that this does not give the length of the entire soleus muscle belly but rather the projection of the instant fascicle length onto the plane of the MTU (Randhawa et al., 2013). The velocity of the muscle fascicles, muscle belly and MTU was then calculated as the first derivative of the lengths over time (shortening denoted with positive sign).

A wireless electromyography (EMG) system (Myon m320RX, Myon AG, Baar, Switzerland, 1000 Hz) was used to measure the surface EMG of the soleus for the same steps as analyzed for the muscle parameters. A fourth-order high-pass Butterworth filter with 50 Hz cut-off frequency, a full-wave rectification and then a low-pass filter with 20 Hz cut-off frequency were applied to the raw EMG data. The EMG values were then normalized for each participant to the maximum obtained during the individual maximum isometric plantar flexion contractions (see below) and averaged over steps. From the normalized EMG curves, we calculated the full width at half maximum (FWHM) as the percentages of stance where the EMG values exceed the half maximum (after subtracting the minimum) (Santuz et al., 2020), to describe the duration of the soleus activation relative to the stance phase.

### Assessment of the soleus muscle intrinsic properties

The soleus force-length relationship was experimentally determined by means of eight maximum voluntary isometric plantar flexion contractions (MVCs) in different joint angles with the right leg on dynamometer (Biodex Medical, Syst. 3, Inc., Shirley, NY) (Bohm et al., 2019). For this purpose, the participants were placed in prone position with the knee flexed to ~120° to restrict the contribution of the bi-articular m. gastrocnemius to the plantar flexion moment (Hof and van den Berg, 1977; Rubenson et al., 2012). The joint angles were equally-distributed from 10° plantar flexion (dynamometer angle) to the individual maximum dorsiflexion angle and set in a randomized order. A standardized warm-up preceded the MVCs. The ultrasound probe and EMG electrodes remained attached between the running and dynamometer measurements.

The resultant moments at the ankle joint were calculated using an inverse dynamics approach under consideration of the effects of gravity and misalignments between joint axis and dynamometer axis (Arampatzis et al., 2005). The respective kinematic data were recorded based on anatomically referenced reflective markers (medial and lateral malleoli and epicondyle, calcaneal tuberosity, second metatarsal and greater trochanter) by means of a Vicon motion capture system (250 Hz). Furthermore, passive moments and the contribution of the antagonistic moments during the plantar flexion MVCs were considered. To assess the contribution of antagonists we used the EMG-based approach reported by Mademli and colleagues (Mademli et al., 2004), under consideration of the force-length dependency of the antagonists (Bohm et al., 2019). The force applied to the Achilles tendon during the plantar flexions was calculated from the joint moment and the individual Achilles tendon lever arm. The ultrasound-based tendon excursion method was used to determine the tendon lever arm (Fath et al., 2010), taking into account its increase during muscle contraction (Maganaris et al., 1998). The soleus muscle fascicle length during the MVCs was synchronously captured by ultrasonography and fascicle length was determined using the aforementioned methodology. Based on the quantified forces and fascicle length in the different ankle joint angles, an individual force-fascicle length relationship was calculated for the soleus by means of a second-order polynomial fit and the maximum muscle force applied to the tendon (F_max_) and optimal fascicle length for force generation (L_0_) was derived, respectively (Bohm et al., 2019; Nikolaidou et al., 2017). The force-velocity relationship of the soleus was further assessed using the dimensionless form (Fung, 1981) of the classical Hill equation (Hill, 1938) and the muscle-specific maximal shortening velocity and the constant a_rel_. For the estimation of the soleus maximum shortening velocity (V_max_) an average fiber type distribution (type 1 fibers: 81.3%, type 2: 18.7%) was assumed based on literature reports (Edgerton et al., 1975; Johnson et al., 1973; Larsson and Moss, 1993; Luden et al., 2008). Using this distribution and reported values of V_max_ for the human soleus type 1 and 2 fibers (Luden et al., 2008) that were adjusted for physiological temperature (37°C) conditions (Ranatunga, 1984), we estimated a V_max_ for the soleus of 6.77 L_0_/s, where L_0_ is the individually measured optimal soleus fascicle length. The constant a_rel_ was calculated as 0.1+0.4FT (Winters and Stark, 1988), where FT is the fast twitch fiber type percentage (a_rel_=0.175). From the force-velocity curve, we calculated the respective power-velocity curve. Based on the soleus force-length, force-velocity and power-velocity relationships, we quantified the individual force-length potential (fraction of the soleus maximum force according to the force–length relationship), force-velocity potential (fraction of the soleus maximum force according to the force–velocity relationship) and power-velocity potential (fraction of the soleus maximum power according to the power–velocity relationship) as a function of the fascicle operating length and velocity during the stance phase of running (Bohm et al., 2019; Bohm et al., 2021a; Bohm et al., 2021b). For the enthalpy efficiency-fascicle velocity relationship we used the efficiency values provided by Hill (Hill, 1964) that were recomputed as a function of shortening velocity according to Barclay (Barclay, 2015). The values were then fitted by means of a cubic spline giving the typical curve with a peak efficiency of 0.446 at a shortening velocity of 0.18 V/V_max_. The resulting function was then used to calculate the enthalpy efficiency of the soleus during running based on the average value of the fascicle velocity over the stance phase (Bohm et al., 2021a; Bohm et al., 2021b).

### Assessment of decoupling within the soleus muscle-tendon unit

The velocity decoupling of the soleus fascicles from the MTU over the time course of the stance phase was quantified as decoupling coefficients. Coefficients were calculated specific for the tendon compliance decoupling (DC_Tendon_, equation 1), fascicle rotation decoupling (DC_Belly_, equation 2) as well as for the overall velocity decoupling of MTU and fascicle velocities that includes both components (DC_MTU_, equation 3) (Bohm et al., 2021b):

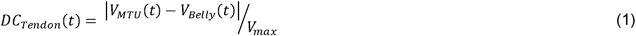

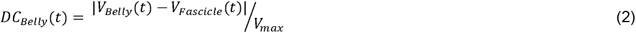

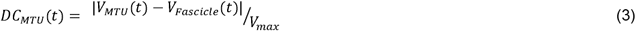

*V(t)* is the MTU, belly and fascicle velocity during the stance phase. These decoupling coefficients were used because previously suggested ratios (i.e. tendon gearing = V_MTU_/V_Belly_, belly gearing (or architectural gear ratio) = V_Belly_/V_Fascicle_, MTU gearing = V_MTU_/V_Fascicle_ (Azizi et al., 2008; Wakeling et al., 2011)) are unfeasible when belly and fascicle velocities become very close to or even zero as during walking and running (Bohm et al., 2018; Bohm et al., 2019).

### Soleus muscle force, work and power estimates

We used a Hill-type muscle model (i.e. muscle force equals the product of the muscle force-length-velocity potential, activation and maximum isometric force) to estimate the normalized soleus muscle forces as a function of time (*t*) during the stance phase using the experimental determined soleus muscle force-length-velocity potentials and muscle activation as inputs:

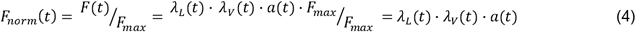

where *F_norm_* is the soleus muscle force normalized to *F_max_, λ_L_* is the force-length potential, *λ_v_* is the force-velocity potential and *a* is the muscle activation. We used the first-order differential equation proposed by Zajac (Zajac, 1989) to estimate the muscle activation from the measured normalized EMG activity *u* (EMG normalized to the maximal EMG activity during the MVCs):

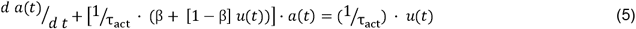

The respective activation time constant (*τ_act_*) and the ratio of the activation to deactivation time constant (*β*) specific to slow and fast twitch fibers were taken from Dick and colleagues (Dick et al., 2017) under consideration of the fiber type distribution mentioned above (i.e. type 1 fibers: 81.3%, type 2: 18.7%). The normalized soleus muscle force was integrated over the normalized fascicle length (normalized to L_0_) to calculate a dimensionless work loop of the soleus during the stance phase of running. The normalized power of the soleus fascicles over the stance phase was calculated as:

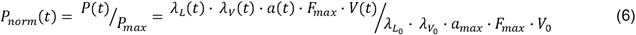

where *P_norm_* is the soleus muscle power normalized to *P_max_, V is* the fascicle velocity, *λ_L_0__* is the forcelength potential at optimal fascicle length, *V_0_* is the optimal fascicle velocity for maximum power, *λ_V_0__* is the force-velocity potential at optimal fascicle velocity and *a_max_* is the maximum activation. *λ*_*L*0_ and *a_max_* are equal to 1.0, thus equation 6 can be rewritten as:

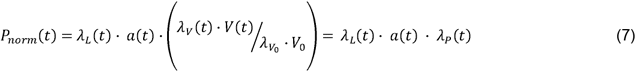

where 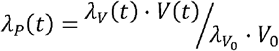

where *λ_P_* is the power-velocity potential.

### Statistics

To investigate the effect of running speed on the outcome parameters, a linear mixed model was applied (factor levels 3 m/s to individual maximal running speed). In case of a main effect of speed, a Benjamini Hochberg corrected post hoc analysis was conducted to control for the false discovery rate and adjusted p-values will be reported. For one participant 8.0 m/s was the highest running speed, thus the maximum speed dataset contained 13 samples (experimental means are reported). In some parameters normal distribution of the normalized residuals was not given (cadence, duty factor, step length, force-length potential, power-velocity potential, efficiency, FWHM, work) as tested by the Shapiro-Wilk test, yet linear mixed models are robust against violations of the normality assumption (Jacqmin-Gadda et al., 2007). The level of significance was set to α = 0.05 and the statistical analyses were performed using R (RStudio v. 2022.07.1, RStudio Inc., MA, USA) and the *nlme* and *emmeans* packages. Unless otherwise indicated, stance phase averaged values will be presented.

## Results

The experimentally determined L_0_ of the soleus was 45.3 ± 6.1 mm and the respective F_max_ was 3486 ± 759 N. The assessed V_max_ of the soleus was 307 ± 41 mm/s. The averaged individual maximum running speed was 8.4 ± 0.2 m/s. There was a significant main effect of speed on stance time, swing time and duty factor (p < 0.001) with decreasing values as speed increased (table 1). Cadence and step length increased significantly with speed (p < 0.001, tab. 1).

**Table 1:** Temporal and spatial gait characteristics at the investigated running speeds (n = 14, mean ± standard deviation).

|  | Running speed |  |  |  |  |  |  |
| --- | --- | --- | --- | --- | --- | --- | --- |
| | 3 m/s | 4 m/s | 5 m/s | 6 m/s | 7 m/s | 8 m/s | 8.4±0.2 m/s |
| Stance time (ms) * | 300±19<sup>a</sup> | 260±17<sup>b</sup> | 223±15<sup>c</sup> | 192±16<sup>d</sup> | 161±11<sup>e</sup> | 138±14<sup>f</sup> | 130±16<sup>g</sup> |
| Swing time (ms) * | 458±27<sup>a</sup> | 444±21<sup>b</sup> | 424±21<sup>c</sup> | 399±15<sup>d</sup> | 371±15<sup>e</sup> | 360±14<sup>e,f</sup> | 354±13<sup>f</sup> |
| Duty factor * | 0.40±0.02<sup>a</sup> | 0.37±0.02<sup>b</sup> | 0.34±0.02<sup>c</sup> | 0.32±0.02<sup>d</sup> | 0.30±0.01<sup>e</sup> | 0.28±0.02<sup>f</sup> | 0.27±0.02<sup>g</sup> |
| Cadence (steps/s) * | 2.68±0.12<sup>a</sup> | 2.88±0.11<sup>b</sup> | 3.10±0.15<sup>c</sup> | 3.44±0.15<sup>d</sup> | 3.80±0.15<sup>e</sup> | 4.02±0.23<sup>f</sup> | 4.15±0.19<sup>g</sup> |
| Step length (m) * | 1.12±0.05<sup>a</sup> | 1.39±0.05<sup>b</sup> | 1.62±0.08<sup>c</sup> | 1.75±0.07<sup>d</sup> | 1.84±0.08<sup>e</sup> | 1.99±0.12<sup>f</sup> | 2.17±0.10<sup>g</sup> |

\* Significant main effect of speed ( $p < 0.05$ ). Values sharing the same letter were not significantly different ( $p > 0.05$ , post hoc analysis).

The soleus MTU lengthened and shortened during the stance phase across all running speeds (fig. 1). The soleus fascicles operated close to optimal length at touchdown and despite the MTU lengthening-shortening behavior, shortened continuously throughout the stance phase at all running speeds (fig. 1). The soleus fascicle shortening showed a significant speed effect (p = 0.011), while post-hoc comparisons demonstrated significantly greater shortening at 4.0 m/s compared to 8.0 m/s and maximum running speed (p = 0.046 and 0.046, tab. 2). The average force-length potential was higher than 0.90 during all speeds and no significant speed-effect was found (p = 0.096, tab. 2, fig. 2). The maximum and average fascicle shortening velocity and EMG activity as well as the FWHM of the EMG activity increased significantly with running speed (p < 0.001, fig. 1, tab. 2). There was a speed-effect on the force-velocity potential with reduced values as running speed increased (p < 0.001, tab. 2, fig. 2). From 3.0 m/s to 6.0 m/s, the average fascicle shortening velocity was very close to the velocity of the efficiency-optimum and, as a result, the averaged enthalpy efficiency was very high (figs. 1 and 3). The higher fascicle shortening velocity at speeds ≥7.0 m/s exceeded the optimum efficiency velocity, which led to a significant reduction of the enthalpy efficiency (p < 0.001, tab. 2). The fascicle shortening velocity at speeds ≥7.0 m/s was, however, very close to the maximum of the power-velocity curve (figs. 1 and 3). Thus, there was a main effect of speed on the power-velocity potential (p < 0.001, tab.2), with higher values at the fast compared to the submaximal running speeds (fig. 3). There was a clear length and velocity decoupling due to tendon compliance and fascicle rotation during the stance phase of running across all speeds (fig. 4). The average values of DC_Tendon_ and DC_Belly_ increased with increasing speed and, as a result, also the overall velocity decoupling between MTU and fascicles, i.e. DC_MTU_ (p < 0.001, fig. 4). Yet, tendon decoupling was always markedly higher than fascicle rotation decoupling (fig. 4).

**Figure 1:**
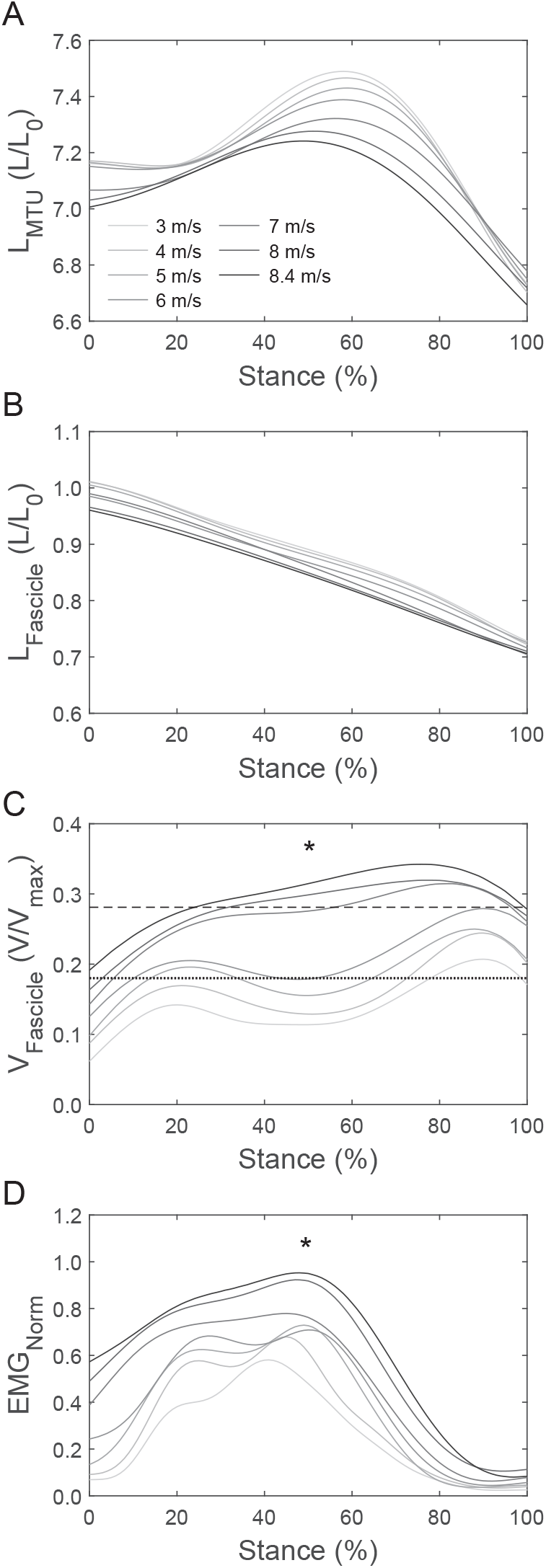
Soleus muscle-tendon unit (MTU, panel A) and fascicle length (L, normalized to optimal fascicle length L_0_, B), fascicle velocity (V, normalized to the maximum shortening velocity V_max_, C) and electromyographic (EMG) activity (normalized to a maximum voluntary isometric contraction, D) during the stance phase of the investigated running speeds. The curves represent the average of the 14 participants and five steps per participant. The dotted line in C indicates the velocity for maximum efficiency (i.e. 0.18 V/V_max_) and the dashed line for maximum power (i.e. 0.28 V/V_max_). * Significant main effect of speed (p < 0.05) on maximum fascicle velocity (post hoc analysis: 3^a^, 4^a,b^, 5^b^, 6^b^, 7^c^, 8^c^, 8.4^c^ m/s, speeds sharing the same letter were not significantly different (p > 0.05)) and maximum normalized EMG activity (post hoc: 3^a^, 4^a,b^, 5^b,c^, 6^b,c^, 7^c,d^, 8.4^d^ m/s), respectively.

**Table 2:**
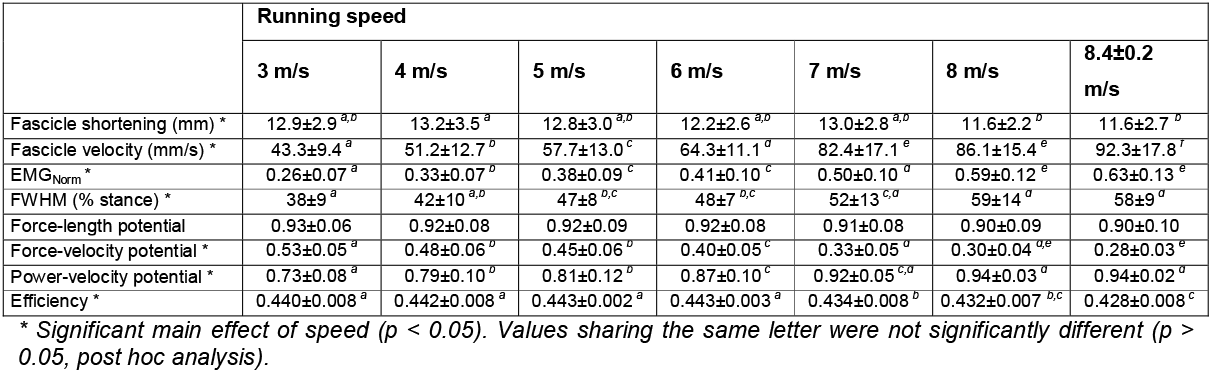
Fascicle shortening, average shortening velocity, average electromyographic activity (EMG, normalized to a maximum voluntary isometric contraction), EMG full width at half maximum (FWHM) and average force-length, force-velocity and power-velocity potential, and enthalpy efficiency during the stance phase of the investigated running speeds (n = 14, mean ± standard deviation).

**Figure 2:**
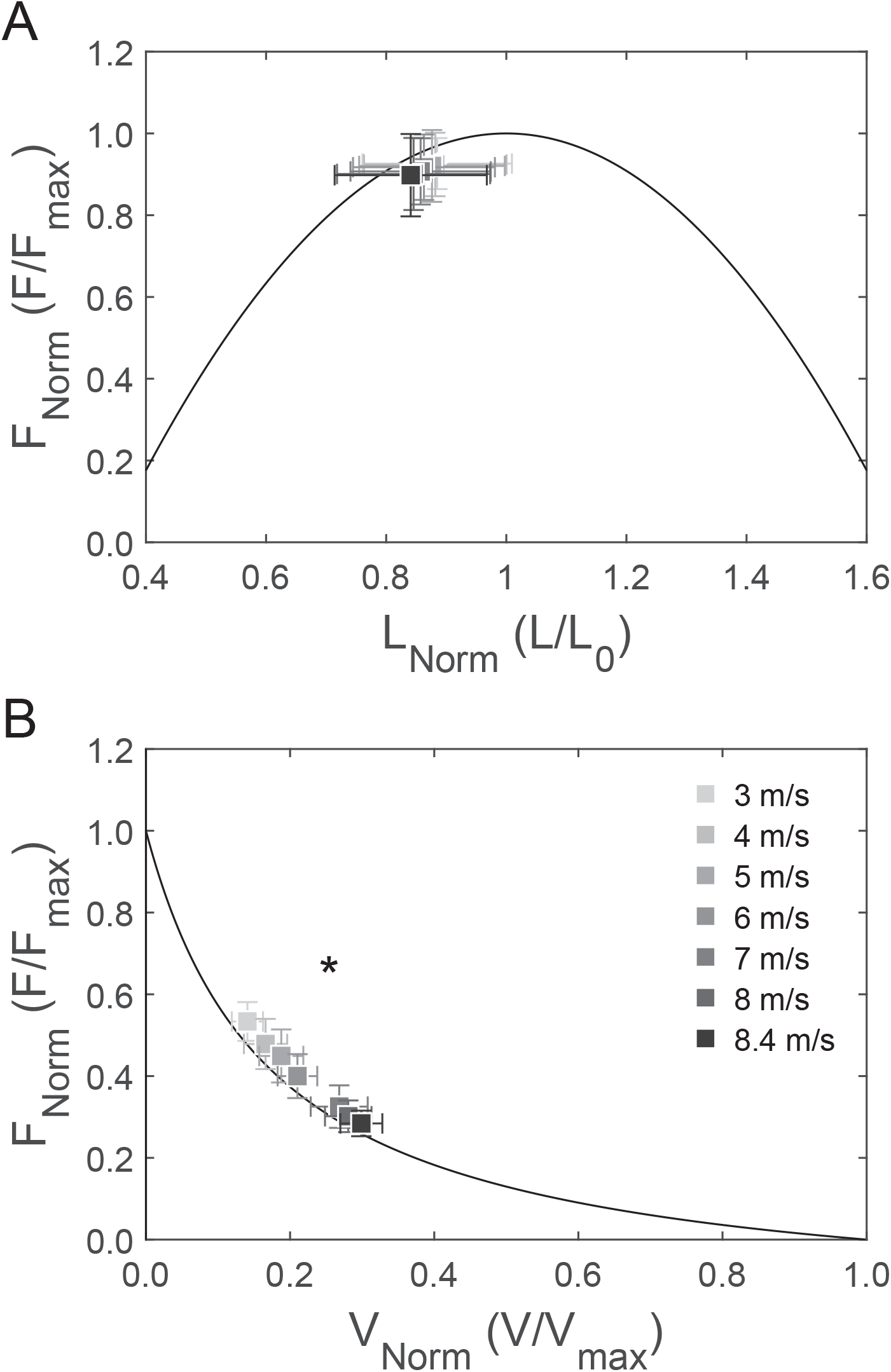
Average operating length (A) and velocity (B) of the soleus fascicles during the stance phase of running at the investigated running speeds mapped onto the normalized force-length and force-velocity curves (n = 14, mean ± standard deviation). Force (F) was normalized to the maximum force (F_max_) during the maximal isometric plantar flexion contractions, fascicle length (L) to the experimentally determined optimal fascicle length (L_0_) and fascicle velocity (V) to the assessed maximum shortening velocity (V_max_). * Significant main effect of speed on force-velocity potential (p < 0.05, post hoc results in table 2).

**Figure 3:**
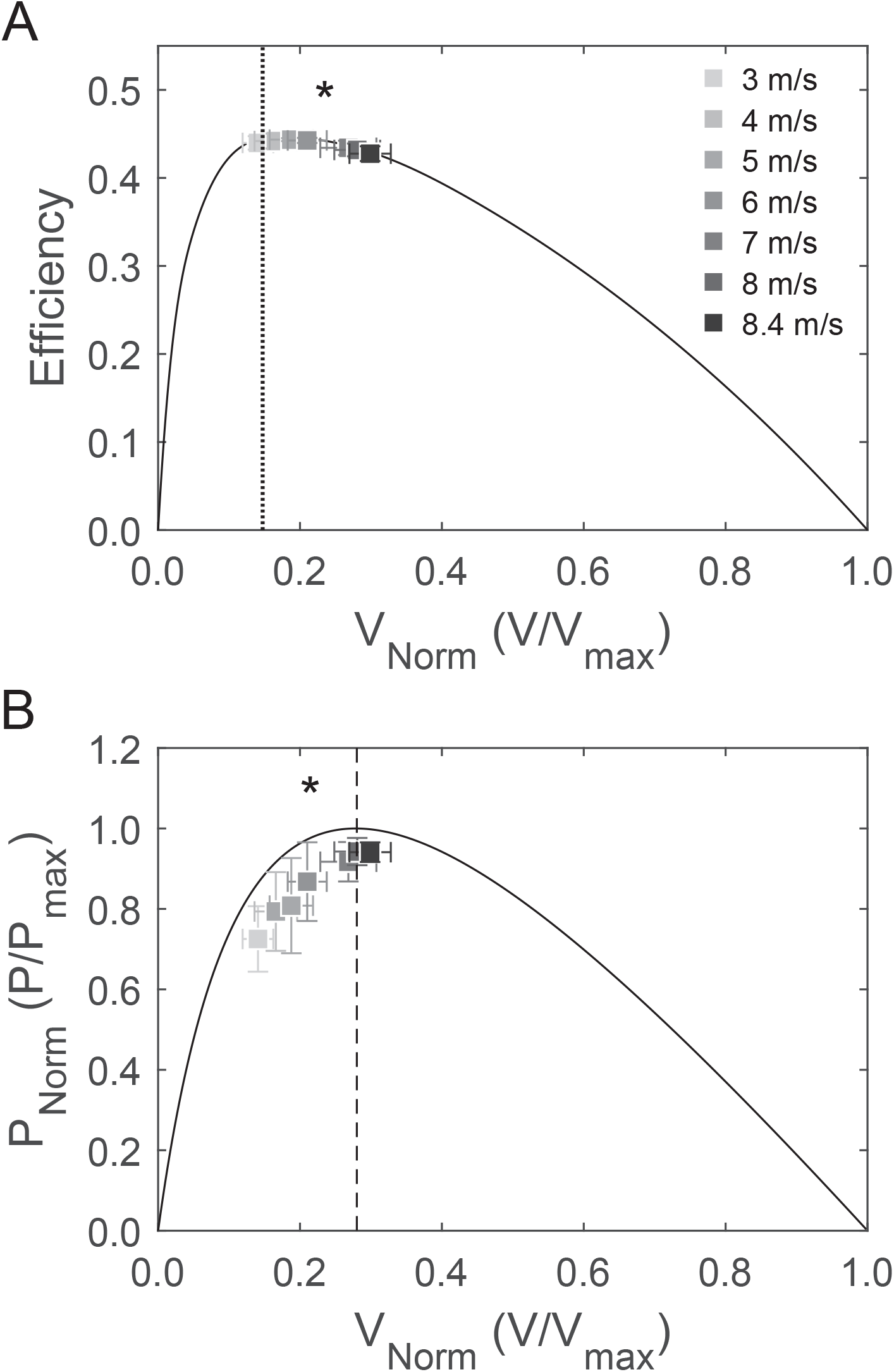
Average operating velocity of the soleus fascicles during the stance phase of running at the investigated running speeds mapped onto the enthalpy efficiency-velocity curve (A) and normalized power-velocity curve (B; n = 14, mean ± standard deviation). Dotted and dashed lines indicate velocities for maximum efficiency (i.e. 0.18 V/V_max_) and power (i.e. 0.28 V/V_max_). Fascicle velocity (V) was normalized to the assessed maximum shortening velocity (V_max_) and power (P) to the maximum power (P_max_) predicted from the force-velocity curve (n = 14, mean ± standard deviation). * Significant main effect of speed on enthalpy efficiency and power-velocity potential (p < 0.05, post hoc results in table 2).

**Figure 4:**
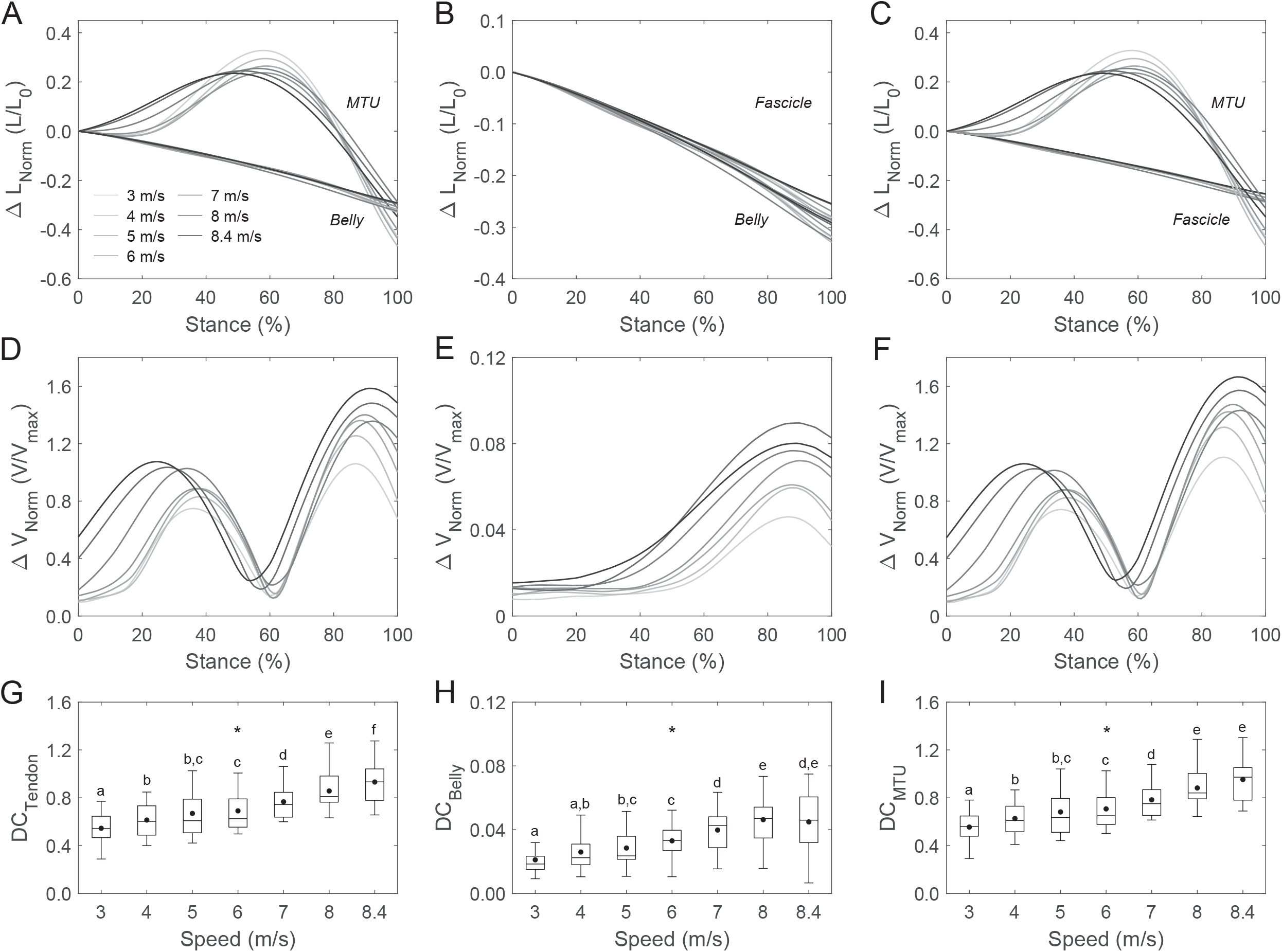
The top row shows soleus muscle-tendon unit (MTU) vs. belly length changes (A), belly vs. fascicle length changes (B), and MTU vs. fascicle length changes (C) normalized to optimal fascicle length (L_0_) during the stance phase of the investigated running speeds. The mid row (D, E, F) shows the resulting velocity decoupling coefficients (DC) and the bottom row (G, H, I) the average DCs (dots indicate the group average). Note the different y-axes scales for the fascicle rotation decoupling. * Significant main effect of speed (p < 0.05), post hoc analysis: box plots sharing the same letter were not significantly different (p > 0.05).

The estimated work loops of the soleus muscle from the Hill-type muscle model featured a counter-clockwise behavior in the muscle force-fascicle length relationship and a continuous production of contractile energy during the entire stance phase in all running speeds (fig. 5). There was a significant speed-effect (p < 0.001) on the normalized maximum soleus muscle force estimates with significantly lower values at the higher speeds (≥7.0 m/s, fig. 5). Although there was also a speed effect (p = 0.030) on the average normalized force during the stance phase, the post hoc comparisons did not reveal any significant differences (p > 0.05) between the different running speeds (3.0 m/s: 0.16±0.03, 4.0 m/s: 0.17±0.04, 5.0 m/s: 0.18±0.05, 6.0 m/s: 0.17±0.05, 7.0 m/s: 0.15±0.03, 8.0 m/s: 0.16±0.03, 8.4 m/s: 0.16±0.03). The produced work by the soleus muscle was similarly distributed between the MTU lengthening and shortening phase and was not affected by speed (p = 0.236, fig. 5).

**Figure 5:**
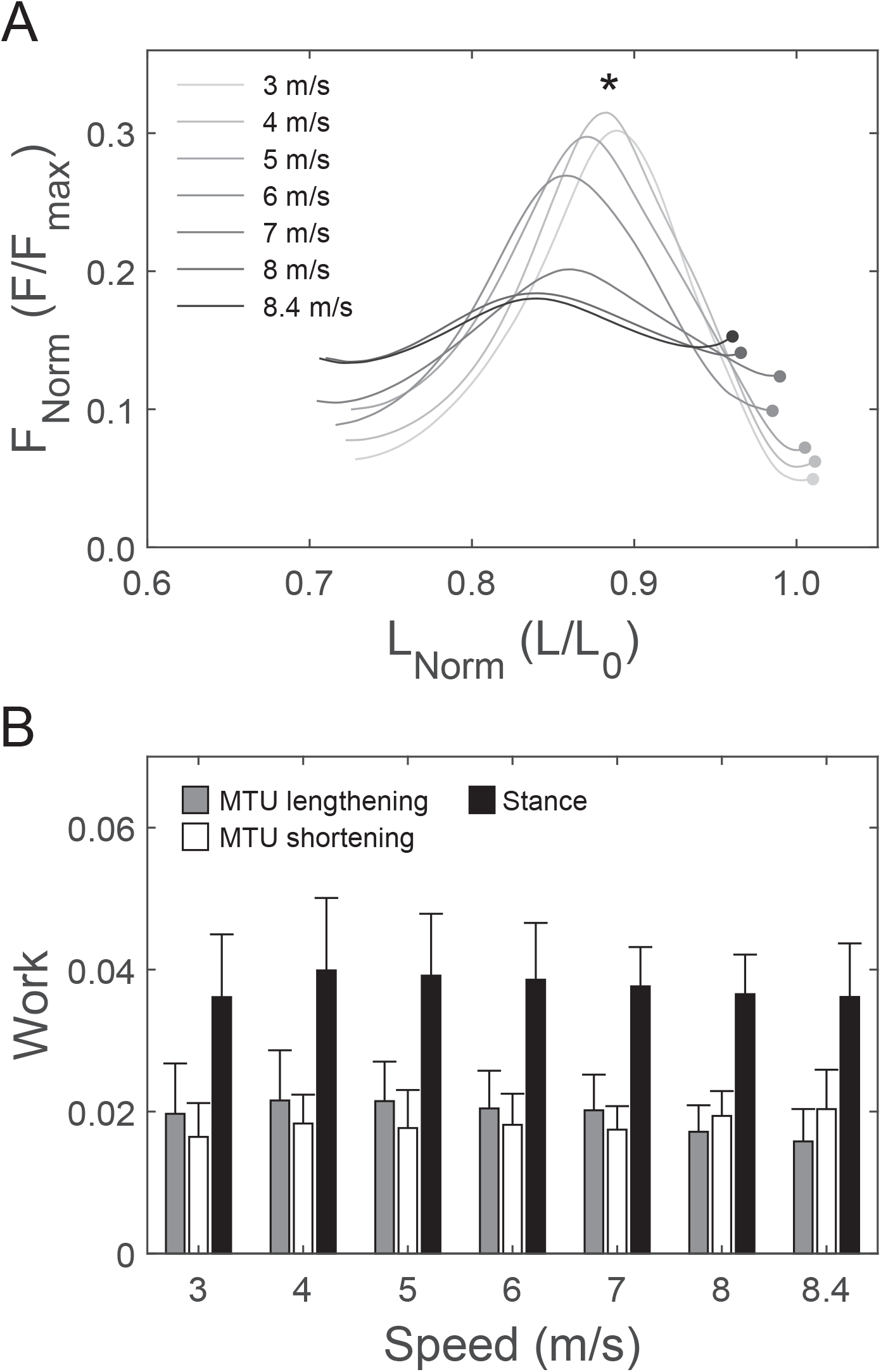
Work loops of the soleus muscle fascicles for the stance phase of the investigated running speeds (A) and produced work (mean ± standard deviation) during the phase of muscle-tendon unit (MTU) lengthening, MTU shortening and over the entire stance phase (B). In A the dots indicate the beginning of the line at touchdown and thus the operating direction during stance (n = 14). * Significant main effect of speed on normalized force maximum (p < 0.05), post hoc analysis: 3^a^, 4^a^, 5^a^, 6^a^, 7^b^, 8^b^, 8.4^b^ m/s, speeds sharing the same letter were not significantly different (p > 0.05).

The contour plot of the normalized soleus muscle force as a function of muscle force-length-velocity potential and activation demonstrated that the decrease in normalized maximum force was due to a pronounced reduction of the force-length-velocity potential (fig. 6A). The increase in maximum muscle activation with speed, although significant (p < 0.001), was not sufficient to compensate for the decline in the force-length-velocity potential (fig. 6A). The contour plot of the estimated normalized soleus power as a function of the force-length-power-velocity potential and activation indicated a contribution of both, activation and contractile conditions to the increase in soleus normalized power until 7.0 m/s speed. At the higher speeds, only muscle activation contributed to the further increase in maximal normalized power (fig. 6B).

**Figure 6:**
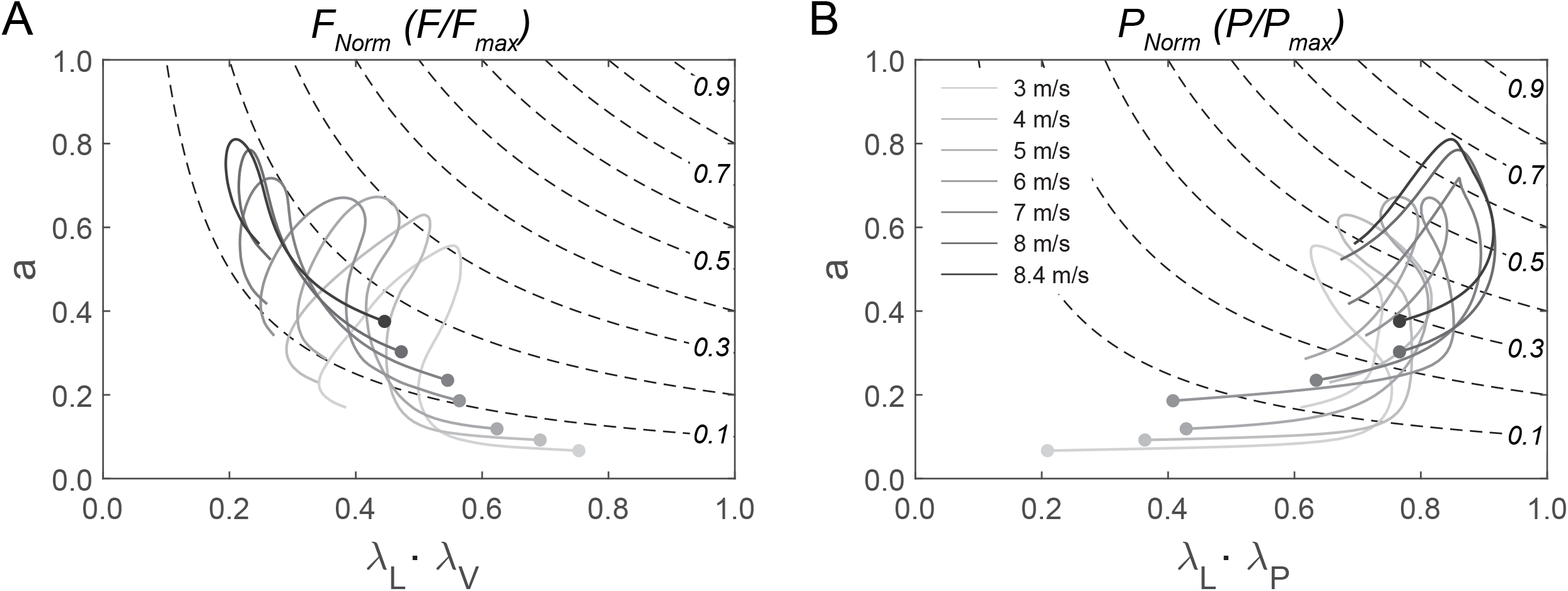
Contour plot for the normalized muscle force (F_Norm_, A) of the soleus as a function of its force-length-velocity potential (λ_L_ * λ_V_) and activation (a). Contour plot for the normalized soleus muscle power (P_Norm_, B) as a function of its force-length-power-velocity potential (λ_L_ * λ_P_) and activation. Respective stance phase trajectories for the investigated running speeds are indicated on top (dots indicate the beginning of the line at touchdown, n = 14).

## Discussion

In the present study, we measured the soleus muscle fascicle behavior from slow to maximum running speed to investigate contractile mechanisms for force generation, power production and efficient mechanical energy production during running. During submaximal speeds (3.0 to 6.0 m/s), the soleus shortened close to optimal length for force generation and at velocities close to the efficiency maximum, two contractile conditions that favor economical muscle work production. With increasing running speed, the soleus continued to operate at fascicle lengths close to the optimal length, yet the fascicle shortening velocity shifted close to the optimal velocity for maximum power production together with a simultaneous increase in the EMG activity. This gives evidence for three neurophysiological mechanisms to enhance muscle power production. Our muscle model estimates predicted a decrease in the soleus maximum force at speeds ≥7.0 m/s as a result of the speed-related impairment of contractile conditions to generate force, particularly the force-velocity potential. Further, the model predicted a continuous increase in the soleus maximum muscle power with running speed, which at speeds >7.0 m/s was mainly due to an increased muscle activation.

With increasing running speed from 3.0 m/s to the individual maximum (i.e. 8.4 m/s), the stance time, swing time and duty factor decreased while cadence and step length increased in magnitudes that are in agreement with earlier studies (Dorn et al., 2012; Santuz et al., 2020). The results showed that during all investigated running speeds, the soleus acted as a work generator by continuous active shortening of its fascicles throughout the entire stance phase (i.e. concentric contraction). This confirms earlier findings from studies on submaximal running (Bohm et al., 2019; Lai et al., 2015; Rubenson et al., 2012; Swinnen et al., 2022), yet gives novel insight into work and power production during maximal running speeds. During the first part of the stance phase at all speeds, the soleus MTU lengthened while the fascicles shortened, indicating a stretch of the Achilles tendon and, thus, storage of elastic strain energy in the tendon. A part of the stored strain energy may originate from a conversion of the bodies kinetic and potential energy, but also the soleus contractile work was stored as elastic strain energy to the tendon. In the second part of the stance phase, where the MTU shortens (i.e. propulsion phase), the soleus muscle continued to produce work. Our model estimates showed that the amount of produced work was not different across all investigated speeds, indicating a consistent energy production by the soleus contractile element during human steady-state running. Furthermore, the produced work was quite similarly distributed between the MTU lengthening and shortening phases, demonstrating that the soleus contractile work done during MTU lengthening is an important fraction of the Achilles tendon energy storage in running.

Active soleus muscle shortening for the required power and work production during running may increase the active muscle volume per unit of force due to the reduced force-velocity potential, which increases the muscle metabolic cost (Roberts, 2002; Swinnen et al., 2023). At the submaximal running speeds (3 to 6 m/s), the soleus fascicles operated at a velocity around the efficiency-optimum and, thus, with a high enthalpy efficiency (>98% of the maximum). In addition, the soleus operated on the upper portion of the ascending limb of the force-length curve very close to the optimal fascicle length. The high average force-length potential (>91%) in turn minimizes the active muscle volume for a given force generation (Beck et al., 2022). The obtained contractile behavior of the soleus muscle indicates an optimization of the tradeoff between the required mechanical energy production and the resulting metabolic energy expenditure during submaximal running, where locomotor economy is arguably a performance criterion (Alexander, 2013; Lazzer et al., 2014). This is in line with recent findings of an improvement in running economy that was associated with a shift of the soleus fascicle operating velocity closer to the optimum of the enthalpy efficiency-velocity curve (Bohm et al., 2021a). Therefore, it is reasonable to argue that the observed almost optimal contractile conditions for work production of the soleus muscle, i.e. high force-length potential and efficiency, minimize the metabolic energy costs during human submaximal running.

At the higher running speeds (≥7.0 m/s), the increased shortening velocity of the soleus fascicles resulted in a decrease of the enthalpy efficiency compared to the submaximal running speeds. Yet, the shortening velocity of the fascicles during the higher running speeds became close to the maximum of the power-velocity curve, increasing the soleus muscle power-velocity potential (i.e. >0.90 at speeds ≥ 7.0 m/s). Considering that also the EMG activity of the soleus muscle was increased with speed and that the soleus force-length potential remained close to its optimum (>0.90), the findings provide first time experimental evidence for a cumulative effect of three different mechanisms, i.e. high muscle force-length potential, high muscle power-velocity potential and high muscle activation, responsible for the required greater soleus power at running speeds above 7.0 m/s. The results further demonstrate that the contractile element of the soleus muscle shifted from favorable conditions to produce mechanical work economically during submaximal running to favorable conditions to produce mechanical power at high to maximal running speeds. Both, tendon compliance and fascicle rotation decoupling mechanisms contributed to the favorable operating length and velocity of the soleus fascicles at the different running speeds and all three decoupling coefficients increased with speed. In the first part of the stance phase throughout all speeds, the compliance of the Achilles tendon allowed the soleus muscle belly to shorten despite lengthening of the MTU. In the second part of the stance phase during MTU shortening, the rotation of the soleus fascicles contributed to the overall MTU decoupling in addition to the tendon compliance. Nevertheless, fascicle rotation decoupling accounted for only 4-5% of the overall MTU decoupling across speeds and, thus, has a comparatively small influence on the soleus fascicle velocities during running.

Using the experimentally determined muscle force-length-velocity potentials and the EMG-derived activation as input variables in the Hill-type muscle model, we predicted a clear decrease in the soleus maximum muscle force at running speeds ≥7.0 m/s despite the increase in muscle activation. The reason was the disproportionately greater reduction of the force-velocity potential compared to the increased muscle activation, indicating that the contractile conditions limit the muscle force generation at maximal running speeds. However, the higher soleus EMG activity with increasing running speed was associated with a widening during the stance phase, i.e. the FWHM increased with speed. This longer duration of activation (relative to the stance phase) resulted in a widening of the estimated work loops of the soleus fascicles (fig. 5A). The result was an almost invariant average soleus muscle force generation and mechanical work production during the stance phase, even though the soleus maximum muscle force substantially decreased at speeds ?7.0 m/s. Recently, Santuz and colleagues reported a widening of the basic activation patterns of muscle groups with increasing running speed using the muscle synergies approach (Santuz et al., 2020). Particularly the temporal component of the propulsion synergy, where the plantar flexors are the main contributors, demonstrated a speed-related widening (Santuz et al., 2020). The widening was interpreted as an increase of the robustness of the neuromotor control to withstand the constrains and challenges imposed by high running speeds (Santuz et al., 2020). The current findings however indicate that the widening of the soleus muscle activation during the stance phase is a relevant neural compensation mechanism to counteract the reduced muscle force potential and thus important for power and work production required for high-speed running. Further, the model predicted a 59% increase of the soleus maximum muscle power from slow to maximum running speed, demonstrating the functional relevance of the power produced by the soleus muscle for increasing running speeds. At speeds until 7.0 m/s, the increase in maximum power was associated with a modulation of both, muscle activation and muscle contractile conditions in terms of its force-length and power-velocity potential. At speeds >7.0 m/s, muscle activation continued to increase, whereas the product of the force-length and power-velocity potential remained quite constant and close to the maximum. This finding suggests that the increase in the soleus maximum power at running speeds above 7.0 m/s was primarily the result of increased muscle activation, while the contractile conditions to produce power were close to their limits. Yet, the increase of muscle power production for high running speeds is a challenge for the metabolic energy supply. The increased soleus muscle activation at the high speeds indicate a greater active muscle volume and, thus, a higher ATP consumption rate (Kipp et al., 2018; Roberts, 2002). In addition, the increased fascicle shortening velocity led to a significant reduction of the enthalpy efficiency at the higher running speeds. While muscle activation increased by 44%, the efficiency decreased by 3% from the slowest to the maximum speed, indicating a more pronounced contribution of the higher active muscle volume, besides the significant effects of the efficiency, to the muscle metabolic energy cost during maximal running.

Non-invasive in vivo investigations of muscle mechanics are challenging and there are some limitations associated with the approach used in the present study that need to be considered when interpreting the results. For the muscle force and power estimates we used a Hill-type muscle model. Although, the experimentally-determined muscle activation and force-length-velocity potentials during running were used as input variables, Hill-type models do not account for the *history dependence of muscle force generation* (Abbott and Aubert, 1952; Rassier and Herzog, 2004). In the present study, the soleus fascicles actively shortened continuously during the stance phase of all running speeds, indicating a shortening-induced force depression that potentially have led to an overestimation of our predicted normalized muscle forces. On the other hand, it has been reported that force depression is independent on muscle shortening velocity but rather depends on the produced muscle mechanical work (Herzog et al., 2000; Kosterina et al., 2008). The predicted mechanical work of the soleus muscle was not affected by running speed, indicating a constant effect of the possible force depression in all running speeds, which would not affect the main findings and conclusions. Further, an average distribution of type 1 and type 2 fibers reported in the literature was used for the assessment of the soleus V_max_, which can differ from the individual values of the investigated participants. In order to examine possible effects of differences in V_max_ on the enthalpy efficiency, force-velocity and power-velocity potentials we performed a sensitivity analysis by changing V_max_ in a substantial range of ±30% in 10% intervals. The sensitivity analysis confirmed the significant main effects of speed, with a decrease in the efficiency and force-velocity potential and an increase in the power-velocity potential with increasing speed (all p < 0.05). Finally, we used the experimentally determined efficiency-velocity curve reported by Hill (Hill, 1964) to relate the measured fascicle velocities to the efficiency. Though the maximum efficiency might differ, the shape of the used enthalpy-velocity curve is very similar to more recent experimental mammalian data (Barclay et al., 2010) and single human slow twitch muscle fibers (He et al., 2000), which indicates minor functional effects on the efficiency results.

In conclusion, our experimental results showed that the human soleus muscle fascicles actively shortened throughout the entire stance phase from slow to maximum running speeds. During the submaximal speeds, the soleus fascicles operated close to the optimum length for force generation with shortening velocities close to the maximum efficiency, thus contractile conditions that promote economical work production. On the other hand, at running speeds ≥7.0 m/s, the shift of the fascicle operating velocity towards the optimum for muscle power production together with the increased muscle activation and the high force-length potential were the three important mechanisms that favored muscle power production. Furthermore, the used Hill-type muscle model suggests a decrease in the maximum soleus muscle force at speeds ≥7.0 m/s as a result of the impaired contractile conditions, i.e. force-velocity potential. Finally, the model predicted an increase in the soleus maximum mechanical power with increasing running speed, mediated by improved contractile conditions and increased muscle activation until 7.0 m/s. At speeds above 7.0 m/s, the contractile capacity of the muscle saturated and the increase in maximum power was driven by an increase in muscle activation.

## List of symbols and abbreviations

EMG: electromyography
MTU: muscle-tendon unit
FWHM: full width at half maximum
F_max_: maximum muscle force
L_0_: optimal length for muscle force generation
V_max_: maximum shortening velocity
FT: fast twitch
DC: decoupling coefficient
MVC: maximum voluntary contraction

## Authors’ contributions

S.B. and A.A. designed research. S.B. performed research and analyzed data. S.B. and A.A. drafted the manuscript. F.M. and A.S. made important intellectual contributions during revision.

## Data availability

The dataset can be accessed from here: 10.6084/m9.figshare.22250104

